# A brain wide circuit model of heat evoked swimming behavior in larval zebrafish

**DOI:** 10.1101/190447

**Authors:** Martin Haesemeyer, Drew N Robson, Jennifer M Li, Alexander F Schier, Florian Engert

## Abstract

Thermosensation provides crucial information but it is poorly understood how temperature representation is transformed from sensation to behavior. Here, we report a preparation that allows control of heat delivery to zebrafish larvae while monitoring motor output and imaging whole-brain calcium signals, thereby uncovering algorithmic and computational rules that couple dynamics of heat modulation, neural activity and swimming behavior. This approach identifies a critical step in the transformation of temperature representation between the sensory trigeminal ganglia and the hindbrain: A simple sustained trigeminal stimulus representation is transformed into a representation of absolute temperature as well as temperature changes in the hindbrain that explains the observed motor output. An activity constrained dynamic circuit model captures the most prominent aspects of these sensori-motor transformations and predicts both behavior and neural activity in response to novel heat stimuli. These findings provide the first algorithmic description of heat processing from sensory input to behavioral output.

## Introduction

Environmental temperature has a strong influence on human behavior, such as seeking shelter or wearing warm clothes in the cold. Similarly, most animal species have a fairly narrow temperature range in which their metabolism can function optimally and evolved behavioral strategies to seek out these preferred temperatures. Especially in cold blooded animals, which cannot actively regulate their body temperature, navigational strategies to avoid extreme heat or extreme cold are of obvious importance. Navigational strategies that lead animals to their preferred temperature within a heat gradient have been studied in diverse species such as E. coli (Maeda et al., 1976), *C. elegans* (Hedgecock and Russell, 1975), zebrafish (Gau et al., 2013) and mouse (Murakami and Kinoshita, 1977).

At the cellular and molecular level it is well understood how animals sense temperature. A large group of transient receptor potential (Trp) channels are gated by temperature, and different Trp channels tile the temperature space from noxious cold to noxious heat (Caterina et al., 1997; Julius and Basbaum, 2001; Schepers and Ringkamp, 2010). In vertebrates, neurons expressing these channels are concentrated in the sensory trigeminal ganglia, where they innervate the face, as well as in the dorsal root ganglia, from where they detect stimuli across the trunk and tail (Erzurumlu et al., 2006; Schepers and Ringkamp, 2010).

Like other sensory stimuli, temperature needs to be encoded and represented by neural activity in primary sensory neurons. This activity then needs to be filtered and processed to extract the information relevant for behavioral responses (Näätänen and Winkler, 1999). Circuit studies have begun to elucidate how the nervous system encodes temperature. Most progress has been made at the periphery. For example, in *C. elegans* the heat sensitive AFD neuron is specifically tuned to changes in temperature via response adaptation (Clark et al., 2006). This strategy is thought to provide information about temperature gradient direction aiding in navigation (Clark et al., 2007). In *Drosophila*, hot and cold sensitive neurons in the antenna form topographic projections in the brain (Gallio et al., 2011), and downstream thermosensory projection neurons which can be subdivided into ON and OFF classes have been implicated in heat avoidance behavior (Frank et al., 2015; Liu et al., 2015). Functional imaging experiments in the mouse have revealed principles of temperature coding in the trigeminal ganglion (Yarmolinsky et al., 2016) as well as the spinal cord (Ran et al., 2016). In the trigeminal ganglion thermosensory neurons tile temperature space, with different neurons responding to different levels of cold or warmth (Yarmolinsky et al., 2016). Overall, most thermosensory neurons represent noxious temperatures while encoding of ambient temperature is sparse (Yarmolinsky et al., 2016). Just like the aforementioned second order projection neurons in *Drosophila*, second order temperature modulated neurons in the mouse spinal cord can be grouped into ON and OFF types (Ran et al., 2016). While warm responsive neurons exclusively show sustained responses, cold-sensitive spinal cord neurons are generally fast adapting (Ran et al., 2016). Despite these advances at the cellular and molecular level, it remains unclear how a temperature percept arises in the brain and how thermosensory activity ultimately leads to behavior. Thus, a description and analysis of the pathways that link temperature sensing to computational processes and behavioral outputs is lacking.

We recently investigated how temperature influences larval zebrafish swimming behavior and found that it is sensitive to both absolute levels as well as changes in temperature and that different behavioral outputs such as turning versus straight swims are differentially influenced by on‐ and off-responsive channels (Haesemeyer et al., 2015). In the current study, we establish brain-wide functional calcium imaging with heat stimulation and behavioral recording to identify heat processing centers throughout the larval zebrafish brain. We find that temperature information is represented in both ON and OFF channels and identify different cell types that represent temperature on different timescales: some are slow-modulated and others fast-adapting. In particular, cell types differ between anatomical regions, with the hindbrain favoring representation on faster timescales while some forebrain regions, including the preoptic area, include cell types that represent temperature on longer timescales. Importantly, we identify a critical step in the sensori-motor transformations: trigeminal sensory neurons represent temperature exclusively using sustained ON and OFF cells, whereas activity diversifies in a trigeminal target area in the hindbrain. There, cell types with transient responses arise and form a more detailed representation of temperature stimulus features. Strikingly, this response type diversification is required to explain the observed behavior. We used these data to derive a realistic circuit model that captures the most important computations underlying the sensori-motor transformations. The circuit model not only captures neural activity transformations but also predicts the behavior and neural activity in response to novel stimuli.

## Results

### A system for brain-wide identification of temperature encoding cells

To observe neuronal activity concurrent with behavior in response to temperature stimuli, we used a fiber-coupled infrared laser to deliver precise heat-stimuli to a head embedded larval zebrafish under a custom-built 2-photon microscope (Figure 1A-B). By keeping the tail of the larva free to move we could monitor behavior under infrared illumination at 100 Hz while simultaneously recording calcium activity at ~2.5 Hz.

**Figure 1.**
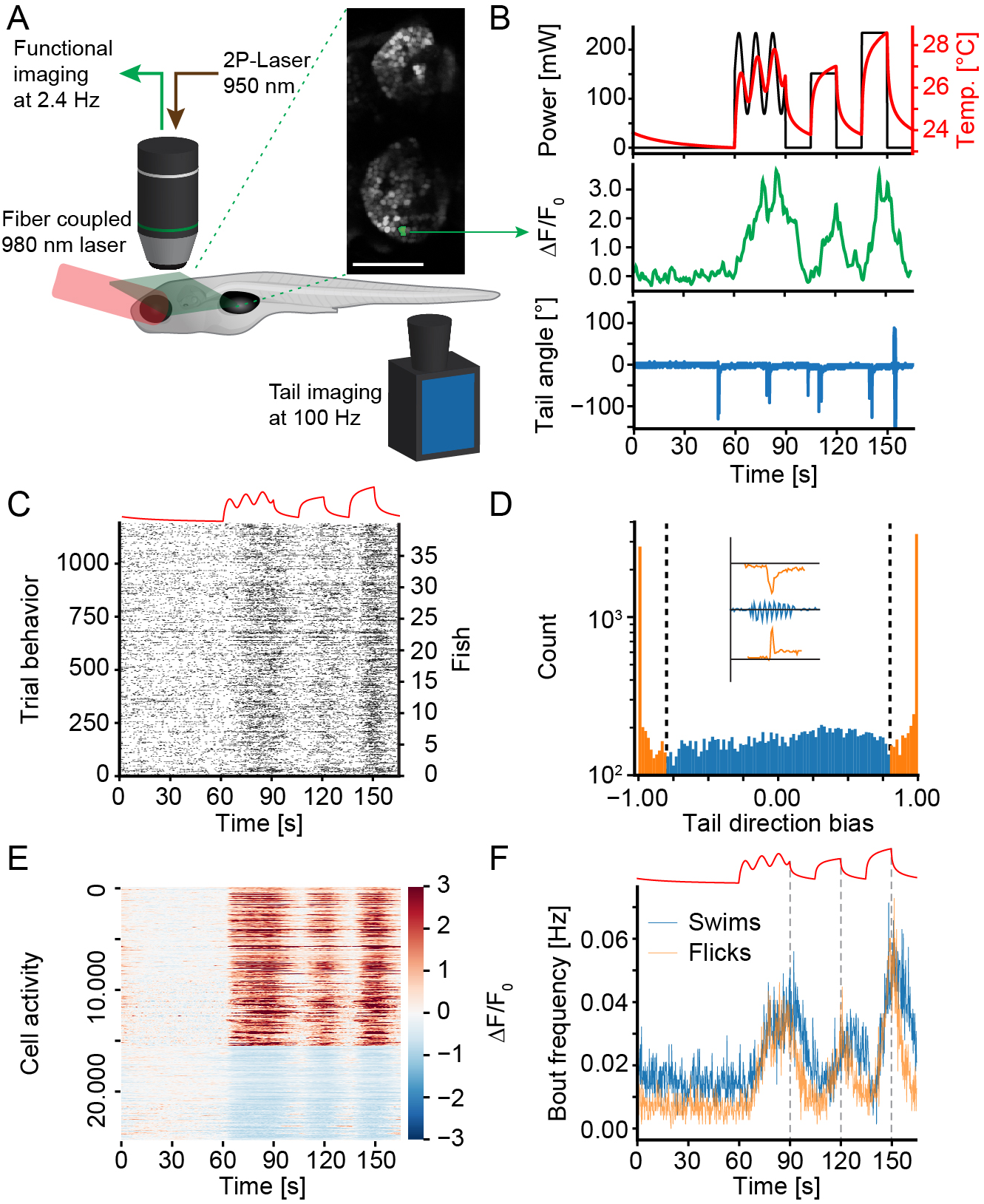
A paradigm to probe heat perception in larval zebrafish. A) Schematic of the setup. Head-embedded tail-free larval zebrafish expressing H2B-GCaMP6s pan-neuronally are imaged at 950 nm under a custom built two-photon microscope. Heat stimuli are delivered using the collimated beam of a fiber-coupled 980 nm diode laser. During imaging, the tail is monitored at 100 Hz to extract behavior. Green plane depicts example imaging plane and inset shows habenulae imaged in one experiment. The activity of the green nucleus is depicted in B. B) Top panel shows the delivered laser power in each repeat (black line) as well as the repeat-average temperature experienced by the fish (red line). Middle panel depicts repeat averaged calcium activity of one example ON cell. Bottom panel depicts example tail-trace during one repeat of one imaging plane. C) Behavior raster plot (summed across repeats) of all 1200 planes imaged across 40 fish. Each black tick identifies the start of a swim bout. Stimulus depicted on top for reference. D) Histogram of directional bias of tail movement across all bouts in all fish. The directional bias is calculated as the sum of angles to the right of the midline during a bout minus the sum of angles to the left of the midline during a bout over the absolute sum of angles. During bouts in which the tail only deflected to the left this will yield a 1, for bouts exclusively to the right a −1 while symmetric tail-undulations will score as 0. Coloring and dashed lines reflect cutoff between “flick” and “swim” categories. Inset shows example tail traces during flick to the right (top), swim (middle) and flick to the left (bottom). E) Heat map of trial averaged activity of all cells across all experiments that have been identified as heat-responsive. Cells are sorted according to ON vs. OFF criteria. Color scale indicates ΔF/F0. F) Experiment average bout frequencies of flick (orange line) and swim (blue line) type bouts. The stimulus is depicted on top for reference. Dashed grey lines indicate start of temperature decline to reveal off response in swims. See also Figure S1.

Since changes in temperature lead to expansion movements in our preparation, we developed an online z-stabilization technique that allowed imaging calcium responses without stimulus induced artefacts (Figure S1A-C). We performed functional imaging experiments in larval zebrafish pan-neuronally expressing the nuclear calcium indicator H2B-GCaMP-6s (Freeman et al., 2014). To study behavioral and neuronal heat responses a simple heat stimulus consisting of both a sinusoidal temperature modulation and discrete temperature steps was presented to the larvae. This stimulus was chosen to probe both sustained and transient heat responses (Figure 1B). With a range of 24 °C to 29°C our heat stimulus stayed below the noxious temperature threshold of ~34 °C. We probed each imaging plane with three trials of the stimulus and imaged 30 planes separated by 2.5 *μ*m in each fish. In total we achieved three-fold coverage of the whole brain across 40 animals.

Across repetitions, the stimulus reliably induced behavior in all tested fish (Figure 1C). As seen in Figure 1B, larval zebrafish do not swim continuously but instead perform swim bouts at discrete intervals (Budick and O’Malley, 2000), and they can elicit bouts of different speeds and turn magnitudes. In our head-embedded larvae data we noticed a prominent class of bouts which were characterized by unilateral flicks of the tail. Therefore using tail dynamics during individual bouts, we subdivided the behavior into undulating “swims” and unilateral “flicks” which likely correspond to near stationary turns in freely swimming behavior (Figure 1D). Importantly, larval zebrafish performed these two behaviors with different dynamics in relation to the temperature stimulus. While both flick‐ and swim-rates rose similarly as temperature increases, swim-rates stayed elevated for a longer time period after the temperature decreased (Figure 1F).

To analyze calcium activity, we anatomically segmented individual cell nuclei. We subsequently used spectral clustering (see Materials and Methods) to extract stimulus evoked activity in an unbiased manner across the whole brain. Imaging a total of 40 fish identified 24,947 responsive cells comprising around 4% of all imaged neurons. The neuronal responses broadly fell into two classes: ON responsive cells, which are excited by increases in temperature and OFF type cells, which are either inhibited by temperature or show rebound excitation when temperature decreases (Figure 1E). Notably, shuffling data with respect to the stimulus reduced the number of identified cells to fewer than 5 % of the original set (Figure S1D). In control fish, which expressed an anatomical indicator (red fluorescent protein, see methods), our clustering approach did not reveal any fluorescence changes resembling our stimulus (Figure S1E-F).

To compare heat induced activity and behavior across stimulus modalities, we imaged a second set of fish presenting a modified heat stimulus followed by an acoustic tap (Figure S1G). These taps elicited escape swims that were in most cases distinct from heat induced unilateral flicks and generally classified as swims, consistent with their bilateral tail dynamics (Lacoste et al., 2015; Figure S1H). In summary, the head embedded preparation enabled characterization of behavioral and neural dynamics across the whole brain while the animal is exposed to temperature and acoustic stimuli.

### Heat related activity is widespread but anatomically clustered throughout the brain

Having established a preparation that allows to monitor temperature modulated neuronal activity, we set out to map heat processing centers throughout the larval zebrafish brain. To this end we registered all acquired imaging data onto a common reference brain (see Methods; Portugues et al., 2014; Rohlfing and Maurer, 2003).

Neurons with heat modulated activity could be identified throughout most of the brain and prominently clustered in specific regions (Figure 2A-D). In the sensory trigeminal ganglia heat sensing neurons occupied a specific caudal subdomain (Figure 2C, right insets). In the hindbrain heat modulated cells formed a cluster in the dorsal cerebellum and another prominent cluster could be identified in rhombomere 5/6 (Rh 5/6) which is a target area for trigeminal fibers carrying information about aversive stimuli (Pan et al., 2012). The forebrain displayed widespread heat related activity with especially large fractions of heat-responsive cells in the sub-pallium as well as the right habenula (Figure 2A-B and E). Furthermore, clusters of heat responsive cells were identified in the pre-optic area which has been implicated in temperature sensation and thermoregulation in mammals (Dean and Boulant, 1989) and reptiles (Cabanac et al., 1967). Brain regions with heat modulated activity generally contained both ON and OFF type cells (Figure 2 C-D), however with varying proportions (Figure 2 E) and with the exception of the cerebellum most regions harbored more ON than OFF type cells.

**Figure 2.**
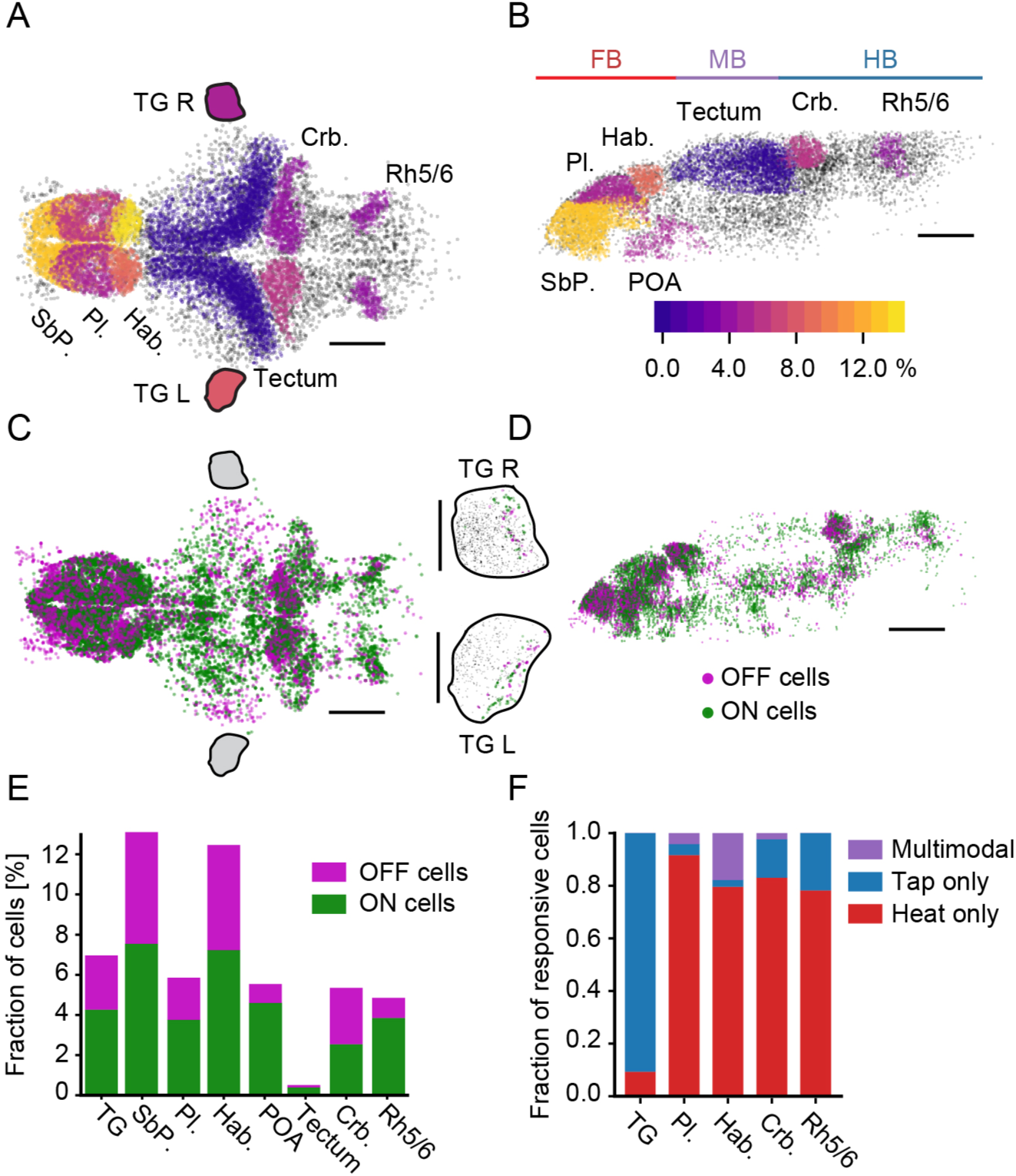
Heat related activity is widespread across the brain. A-B) Fraction of heat responsive cells within selected brain regions. Color scale indicates percentage of heat-sensitive cells within each selected region. Grey cells were not analyzed for this panel. Scale bars, 100*μ*m Pl.: Pallium, SbP.: Subpallium, Hab.: Habenula, Crb.: Cerebellum, Rh5/6: rhombomeres five and six of the hindbrain, POA: Preoptic area, TG L: Left trigeminal ganglion, TG R: Right trigeminal ganglion. Colored lines on top delineate major subdivisions of the brain, FB: Forebrain, MB: Midbrain, HB: Hindbrain. Note that trigeminal ganglia are not to scale. A) Dorsal view of the brain, anterior left, left side bottom. B) Side-view of left hemisphere, anterior left, dorsal top. C) Distribution of ON (green) and OFF (magenta) cells across the zebrafish brain (top projection). The projection shows all cells identified across 30 individual experiments which have been registered onto a common reference brain. Scale bar 100 *μ*m, anterior left, left side bottom. Black outlines mark approximate location of trigeminal ganglia which are shown in insets to the right (TG R right trigeminal ganglion, TG L left trigeminal ganglion). Each trigeminal ganglion depicts cells across five fish registered onto a common reference ganglion. Scale bar 50 *μ*m, anterior left. D) Side-view of the brain in C), only cells in the left hemisphere are depicted. Scale bar 100 *μ*m, anterior left, dorsal top. E) Fraction of heat ON cells (green) and heat OFF cells (magenta) in select brain regions. F) For regions that were imaged in heat and tap experiments depicts the fraction of stimulus responsive cells that only responded to the heat stimulus (red), cells that only responded to the tap stimulus (blue) and multimodal cells that responded to both heat and tap (purple). See also Figure S2.

After the identification of heat processing centers, we tested whether heat-modulated cells and their ON and OFF subtypes cluster in the brain or if they are rather distributed in a random manner. Comparing nearest neighbor distances within and across types revealed that heat modulated cells indeed clustered together and this was maintained across specific subtypes such as ON and OFF cells as well (Figure S2A). This clustering indicates that functional subdivisions are reflected in the anatomical location of cell types. Notably, shuffling cell identities removed all region-specific cell and type clustering (Figure S2B-D) and abolished differences in nearest neighbor distance metrics (Figure S2E). This confirms that the observed structure in the data is indeed a feature of the brain and does not simply arise by chance.

After the anatomical characterization we wanted to know whether heat modulated neurons likely encode heat information specifically or if some neurons generalize across stimulus modalities. To distinguish between these possibilities, we used the stimulus set combining temperature changes and aversive acoustic taps (Figure S1G) and identified cells that only respond to either the temperature or tap stimulus alone (unimodal cells) as well as cells that respond to both (multimodal cells) (Figure S2F). Importantly, the sensory trigeminal ganglia contained only unimodal cells for tap or for heat (Figure 2F blue and red) a property which was also largely reflected by cells in the hindbrain (Figure 2F). While a much larger fraction of trigeminal cells responded to the strong tap stimulus rather than temperature this distribution reversed in the hindbrain. Indeed, mechanosensory *islet1* expressing cells in the trigeminal do not form extensive arborizations in rhombomeres 5/6 of the hindbrain which could explain this observed difference (Pan et al., 2012). The forebrain on the other hand contained a significant fraction of multimodal cells and taps were largely represented by these. Especially in the habenula, tap responsive cells were almost exclusively multimodal, which suggests that taps are not encoded there with independent negative valence (Figure 2F). In summary, the data demonstrate that heat evoked activity is widespread throughout the brain but heat responsive neurons nonetheless cluster into specific regions such as the posterior trigeminal ganglion, rhombomeres five and six of the hindbrain or the cerebellum. Furthermore, while most neurons seem to be modality specific, especially in the forebrain cell types arise that have a mixed representation of aversive stimuli.

### Motor cells encode swim types and are stimulus dependent

After pinpointing neurons and brain regions processing temperature stimuli we next sought to identify neurons with motor-correlated activity. To this end we used the bout starts in each imaging plane (Figure 1C) to derive behavioral regressors by convolution with a calcium response kernel (Miri et al., 2011). These regressors represent the expectation of calcium responses in a cell that encodes the behavior and can therefore be used to probe the brain for cells that show activity which is strongly correlated (*r* ≤ 0.6) to motor output (Figure 3A).

**Figure 3.**
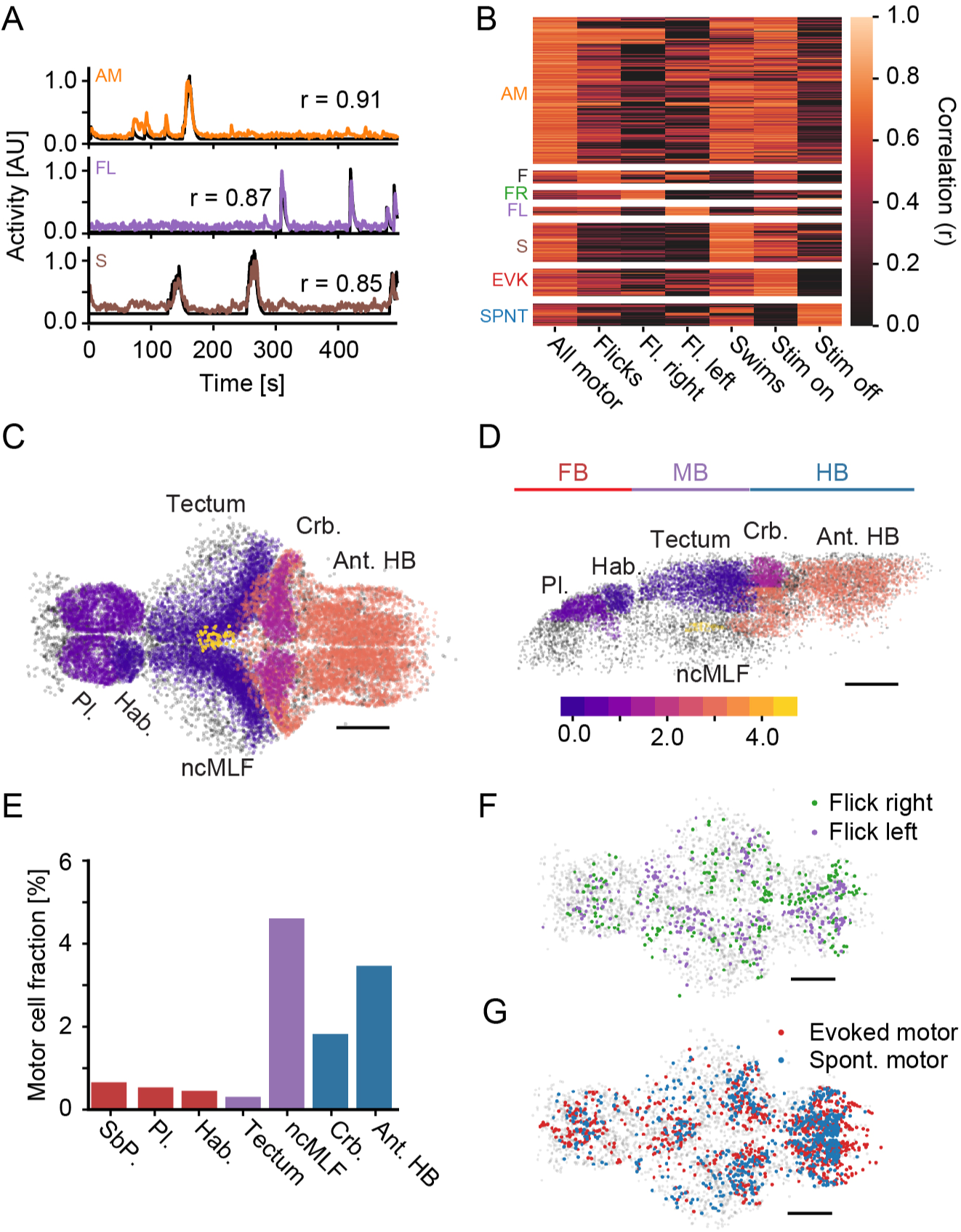
Motor cells can be separated according to behavior and stimulus conditions. A) Example behavioral regressors (black) and activity trace of one correlated cell. Top: Cell encoding all motor events in a plane (orange); Middle: Cell encoding left flicks in a plane (purple); Bottom: Cell encoding swims in a plane (brown). Numbers indicate correlation coefficient. B) Clustered heatmap of correlations of motor-cell activity to individual motor regressors. All motor: Regressor included all motor events of a given plane. Flicks: Regressor only included flicks. Right flicks: Regressor only included right flicks. Left flicks: Regressor only included left flicks. Swims: Regressor only included swims. Evoked motor: Regressor only included motor events while the heat stimulus was on. Spontaneous motor: Regressor only included motor events while the heat stimulus was off. Cells are only assigned to a more specialized motor cluster if the correlation to the specialized regressor is significantly higher than to the general regressor (*p* < 0.01, bootstrap hypothesis test). All-motor (AM): N = 5049 cells; Flicks (F): N = 420; Flick-right (FR): N = 319; Flick-left (FL): N = 298; Swims (S): N = 1338; Evoked-motor (EVK): N = 950; Spontaneous-motor (SPNT): N = 763 C-D) Fraction of motor correlated cells within selected brain regions. Color scale indicates percentage of motor-correlated cells within each identified region. Scale bars, 100*μ*m. Pl.: Pallium, Hab.: Habenula, ncMLF: nucleus of the medial longitudinal fascicle, Crb.: Cerebellum, Ant. HB: Anterior hindbrain. Grey cells were not analyzed for this panel. C) Dorsal view of the brain, anterior left, left side bottom. D) Side-view of left hemisphere, anterior left, dorsal top. E) Quantification of percentage of motor correlated cells in select brain regions. Red bars: Forebrain; purple: Midbrain; blue: Hindbrain. F) Distribution of Flick-right (green) and Flick-left (purple) cells, top projection. Anterior left, scale bar = 100 *μ*m G) Distribution of Evoked-motor (red) and Spontaneous-motor (blue) cells, top projection. Anterior left, scale bar = 100 *μ*m. Grey cells in F-G are non-motor related cells representing brain outline. See also Figure S3.

We generated regressors encoding all motor events (Figure 3A, “AM”) as well as regressors encoding the two subsets of motor events, flicks (“FL”) or swims (“S”). Correlating activity across the whole brain to these regressors revealed a large representation of all motor events as well as neurons significantly more correlated to regressors encoding behavioral submodules such as flicks to the right and left or swims (Figure 3B, *p* < 0.01 bootstrap hypothesis test).

Motor events could either be controlled by a single set of pre-motor cells or different stimulus modalities could influence different pre-motor pools. We therefore probed the brain for cells that encoded motor events in a stimulus dependent manner. Namely, we created regressors that only reported motor output while the stimulus is being delivered or conversely during rest, while the stimulus is off. These regressors could indeed recover cells that encode behavior contingent on the stimulus presentation, “Evoked-” and “Spontaneous-motor” cells (Figure 3B, “EVK” and “SPNT” respectively). Bout triggered averages revealed the specific responses of motor cell types during bouts. As expected, while “All-motor” cells were equally responsive during left and right flicks, “Flick-left” and “Flick-right” cells responded almost exclusively during left and right flicks, respectively (Figure S3A). The bout triggered averages also revealed that “All-motor” cells respond with equal strength irrespective of the stimulus, while “Evoked-motor” and “Spontaneous-motor” cells showed a much stronger response in the presence or absence of the stimulus, respectively (Figure S3B). Importantly, shuffling the activity data with respect to the behavior reduced the number of identified cells to less than 1.8% (Figure S3C) and removed all structure from the bout triggered averages (Figure S3D).

The anterior hindbrain and cerebellum contained prominent clusters of motor related cells (Figure 3C-D and Figure S3E-F). We also identified a concentration of motor encoding cells in the nucleus of the medial longitudinal fascicle (ncMLF), which has been implicated in controlling swim speed (Severi et al., 2014; Figure 3C-D). While a sizeable fraction of hindbrain and ncMLF neurons encoded motor behavior, there were only few such neurons in the forebrain (Figure 3E). Cells encoding flicks to the left versus right were notably absent from the ncMLF and showed a lateralized distribution, especially in the hindbrain, where more cells encoded behavior in an ipsilateral manner (Figure 3F). Evoked‐ and Spontaneous-motor cells on the other hand were largely dispersed throughout motor related brain regions but did cluster spatially within those regions (Figure 3G).

In summary, behavioral subtypes and stimulus contingencies are encoded by separate pools of motor cells that are largely confined to regions previously described as encoding motor activity (Dunn et al., 2016; Naumann et al., 2016; Portugues et al., 2014).

### Activity decorrelation in the hindbrain is required to explain behavior

After the identification of heat processing centers and motor related cells in the brain we wanted to know how well temperature related activity in specific regions can explain the observed swim and flick behaviors. To this end we partially annotated our reference brain using Z-Brain annotations (Randlett et al., 2015). This allowed extracting sensory related activity using spectral clustering for cells in specific regions, effectively capturing more diversity than brain-wide clustering (see Materials and Methods). This analysis revealed that stimulus representation in the sensory trigeminal neurons is simple, consisting of one ON and one OFF cell type tracking the stimulus with slow dynamics (Figure 4A). Furthermore, the activity of both cell types was highly anti-correlated (Figure 4A’), indicating that they have very similar stimulus encoding albeit with opposite sign.

**Figure 4.**
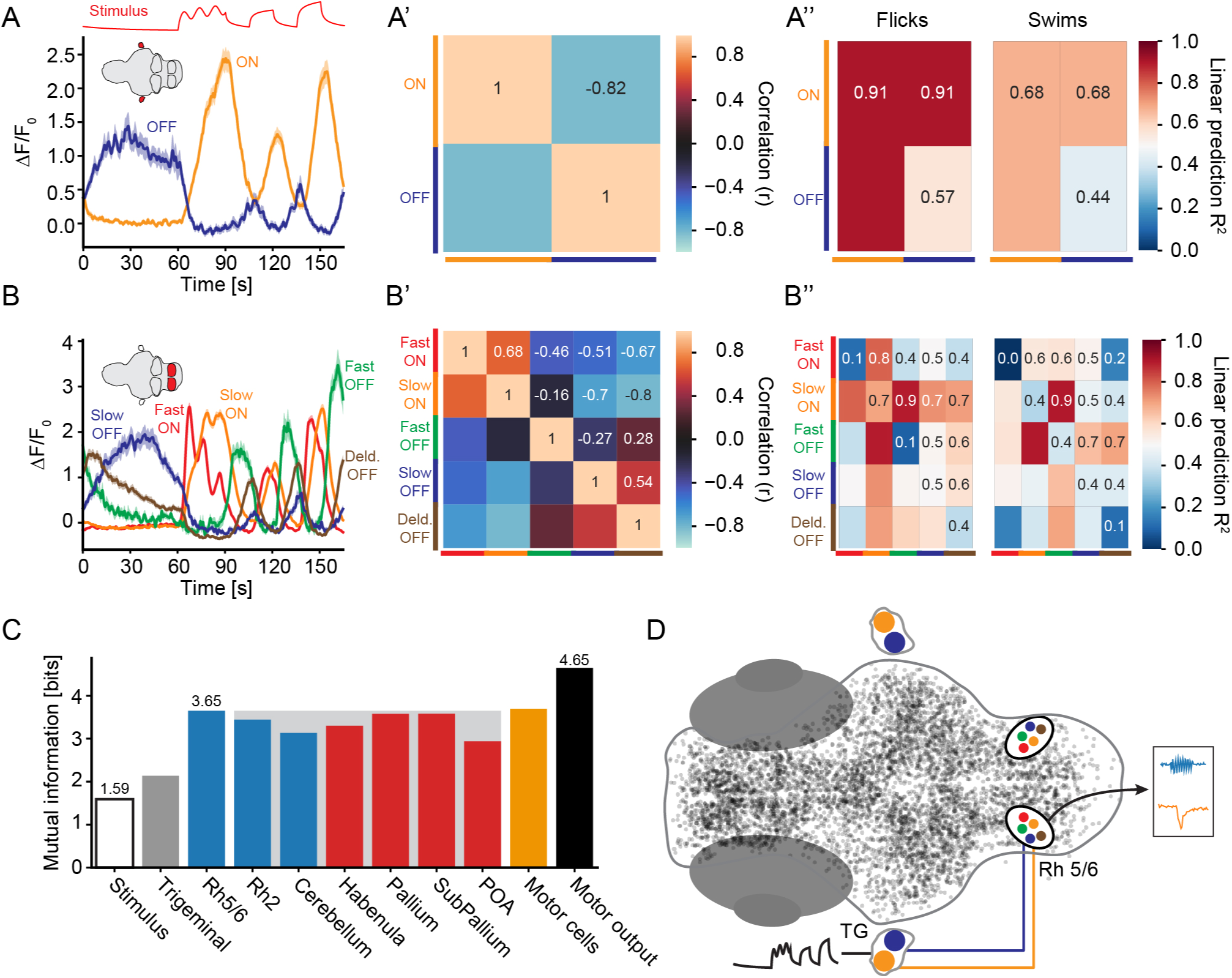
Diversity of heat responses increases in the hindbrain. A-A”) Characterization of heat responses in the trigeminal ganglion. A) Response types extracted via spectral clustering, ON cells orange, OFF cells blue. Thick lines indicate cell-average activity, shading indicates bootstrap standard error. A’) Pairwise correlations of the response types to quantify similarity. A”) Coefficient of determination (*R*^2^) for using one (diagonal) or up to two of the response types to predict flicks (left panel) or swims (right panel). B-B”) Characterization of heat responses in the Rhombomere 5/6 region of the hindbrain B) Response types extracted via spectral clustering. Fast-ON cells red, Slow-ON cells orange, Fast-OFF cells green, Slow-OFF cells blue and Delayed-OFF cells brown. Thick lines indicate cell-average activity, shading indicates bootstrap standard error. B’) Pairwise correlations of the response types to quantify similarity. B”) Coefficient of determination (*R*^2^) for using one (diagonal) or up to two of the response types to predict flicks (left panel) or swims (right panel). A linear model combining just two activity types was chosen for direct comparison with the trigeminal activity types. C) Mutual information with the motor output by knowing the stimulus (hollow bar) or all heat-related activity in the given brain regions (grey trigeminal ganglion, blue hindbrain, red forebrain) or motor cell activity (orange bar). The filled black bar quantifies the entropy in the motor output itself. The height of the grey box indicates mutual information in Rh 5/6 and marks regions not included in the circuit model. D) Schematic of response diversification between detection in the trigeminal and cells in rhombomeres 5 and 6 of the hindbrain followed by the generation of motor output (black arrow). Colors indicate response types. See also Figure S4.

We next asked whether a simple linear regression model using the activity present in the trigeminal sensory ganglion could explain the observed flick and swim rates. Activity in the trigeminal was indeed sufficient to explain flick type bout generation, capturing more than 90 % of the variance in this behavior (Figure 4A” left) but was considerably worse in explaining swims, capturing less than 70% of the variance (Figure 4A” right).

In the rhombomere five and six region of the hindbrain, a trigeminal target area (Pan et al., 2012), activity profiles became considerably more diverse with the presence of two distinct ON and three distinct OFF cell types (Figure 4B). Importantly, both transient “Fast-ON” and transient “Fast-OFF” activity arose in this region while another set of “Slow-ON” and “Slow-OFF” neurons mostly reflected trigeminal activity (Figure 4B). These new response types change the stimulus representation from exclusively encoding temperature levels to also representing the direction of temperature change. The distinct temporal dynamics furthermore resulted in a decorrelation of activity at this first relay station (Figure 4B’). Remarkably, combining the activity of just two response types, the trigeminal-like Slow-ON and a newly formed Fast-OFF type was sufficient to explain both flicks and swim behavior, capturing 90% of the variance in both behaviors (Figure 4B”). This suggests that the observed response diversification underlies the generation of behavior.

Analyzing activity in other brain regions revealed differences in stimulus representation throughout the brain (Figure S4A-E). Temperature related activity in the cerebellum followed different dynamics than activity in Rh 5/6, and the representation of stimulus amplitude was strongly reduced in this region (Figure S4A). The forebrain, especially the pallium (Figure S4C) and preoptic area (Figure S4E), contained activity types that evolve on considerably slower timescales than displayed by neurons within the hindbrain (Figure 4B, Figure S4A). This might indicate a role of forebrain areas in longer timescale integration of stimuli potentially related to the setting of behavioral states rather than generation of short-timescale behavior.

To describe the amount of information region specific activity carries about the observed behavior, we analyzed mutual information between activity in each region and the flick and swim behavior rates. Mutual information between all activity in a given region and the observed motor output reflected the results of the linear model (Figure 4C). While there was a modest increase in mutual information between considering only the sensory stimulus or activity in the trigeminal ganglion, activity in rhombomeres 5/6 of the hindbrain contained as much information about behavior as activity in any other region, except for the motor cells themselves (Figure 4C). Therefore our preferred model is that other brain areas like the pallium and subpallium or cerebellum are not directly involved in the described behavior but rather monitor the information for higher order purposes. Shuffling the activity data as a control reduces the number of cells identified through clustering to less than 3% in each region (Figure S4F). This argues that the recovered cell types are a true feature of stimulus representation.

In summary, the data indicate that activity transformation in a trigeminal target area (Figure 4D) is an important step in the observed sensori-motor transformations: this transformation is necessary to explain the behavioral output while later stimulus transformations do not seem to increase information about the motor output (Figure 4C).

### A dynamic circuit model captures activity transformations and generation of behavior

To better understand and describe the sensori-motor transformations underlying heat evoked swimming behavior we sought to build a dynamic circuit model that is constrained by the behavior and the observed neuronal activity. This model describes four independent, sequential transformations: First, sensory stimulus to activity in the trigeminal; second, activity in the trigeminal to activity in Rh 5/6; third the transformation from Rh 5/6 to Motor cells, and lastly the generation of behavior given the activity in the Motor cells (Figure 5).

**Figure 5.**
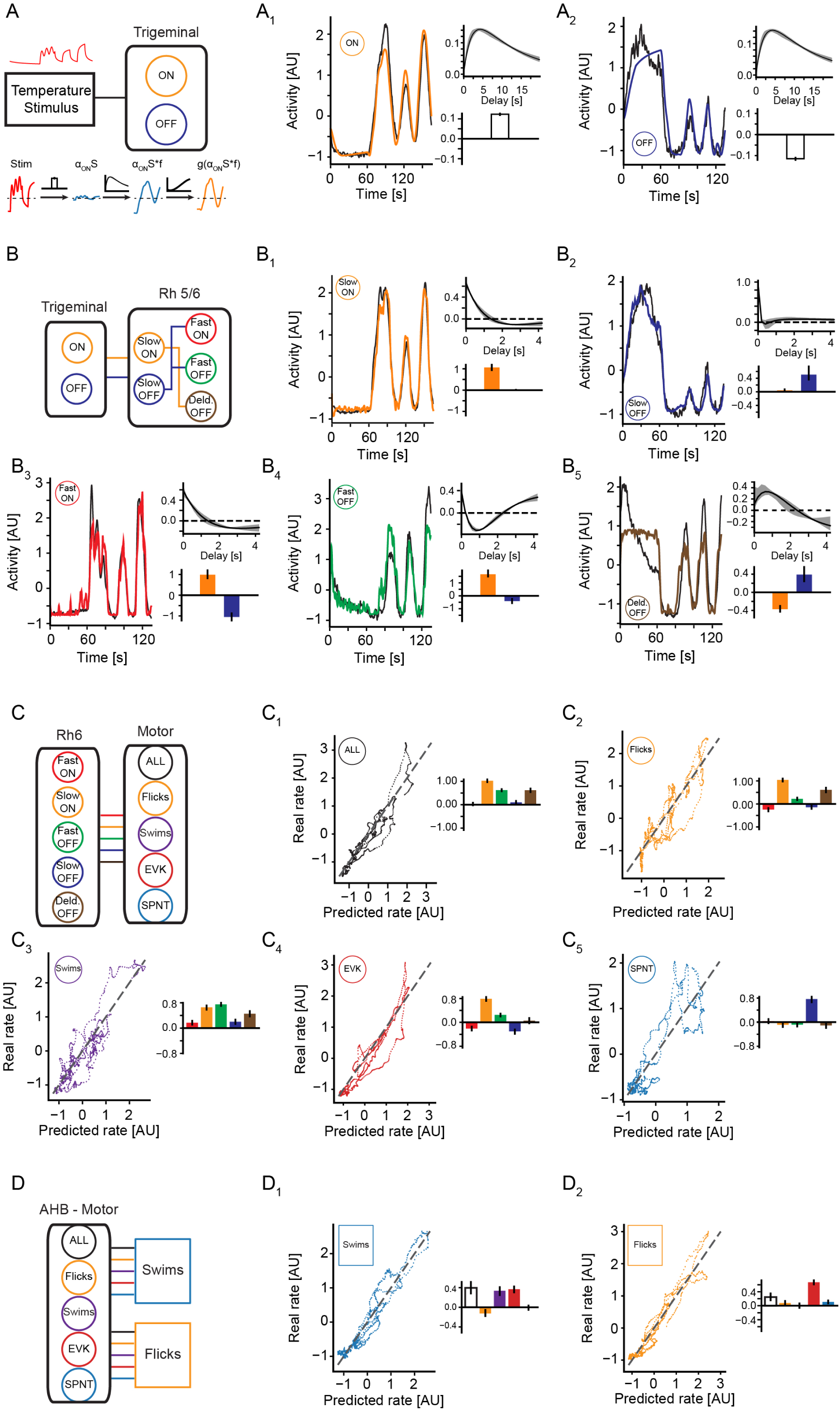
A dynamic model of sensori-motor transformation during heat perception. A) Schematic of the first model stage which relates sensory heat input to activity in the two trigeminal cell types. Red curve depicts sensory stimulus of experiments used for fitting the model. Bottom panel is schematic depiction of the influences of the individual components (linear factors, filter, nonlinearity) of the dynamic model using the trigeminal ON cell type as an example. A_1_) Model prediction of trigeminal ON activity (orange) and measured activity (black) (left panel), impulse response of the model filter (top right) and linear coefficient (bottom right). A_2_) Model prediction of trigeminal OFF activity (blue) and measured activity (black) (left panel), impulse response of the model filter (top right) and linear coefficient (bottom right) B) Schematic of the second model stage which relates trigeminal output activity to the activity types observed in Rhombomeres 5/6 of the hindbrain. Note that the three types in the right column rely on indirect inhibition via the slow ON or slow OFF types. B_1_) Model prediction of Slow-ON activity (orange) and measured activity (black) (left panel), impulse response of the model filter (top right) and the linear coefficients for the trigeminal ON (orange) and OFF (blue) cells (bottom right). B_2_) - B_5_) Same as B_1_) but for Slow-OFF (B_2_), Fast-ON (B_3_), Fast-OFF (B_4_), and Delayed-OFF (B_5_) types. C) Schematic of the third model stage relating output rates in the hindbrain units to activation rates of the motor correlated cells C_1_) Scatter plots of actual versus predicted output rates of modeled All-motor activity (left panel) and the linear coefficients for the Fast-ON (red), Slow-ON (orange), Fast-OFF (green), Slow-OFF (blue) and Delayed-OFF (brown) types. C_2_) - C_5_) Same as C_1_) but for Flicks (C_2_), Swims (C_3_), Evoked-Motor (C_4_) and Spontaneous-Motor (C_5_) types. D) Schematic of the last model stage relating output rates of the motor correlated cells to behavioral rates. D_1_) Scatter plots of actual versus predicted behavior rates of modeled swim output (left panel) and the linear coefficients for the All-motor (black), Flick (orange), Swims (purple), Evoked-motor (red) and Spontaneous-motor (blue) cells. D_2_) Same as D_1_ but for predicted behavior rates of flick output. Shading and error bars indicate 99 % confidence intervals after sampling from the posterior distribution. See also Figure S5.

The transformation from sensory stimulus to trigeminal activity is a dynamic process and relies on temporal processing. We therefore fit a model that combines a linear multiplication of the sensory input with convolution by a temporal filter (Figure 5A). Both the multiplicative term and filter parameters were fit using Markov-chain-Monte-Carlo sampling, obtaining confidence intervals on the parameters in the process (Hastings, 1970; see Materials and Methods for details). Since stimulus encoding relied on a non-linear transformation, a cubic nonlinearity accounted for differences in mapping between stimulus strength and neuronal activity (Figure S5 A-B). This approach allowed explaining the observed ON and OFF type activity in the trigeminal ganglion in terms of the sensory stimulus as evidenced by the close juxtaposition of the fits and true activity (Figure 5A_1_-A_2_). As expected, the linear factors of the model demonstrate an activation of the ON type and an inhibition of the OFF type by the sensory stimulus. As expected at this stage a linear filter resembling the kernel of a nuclear calcium indicator (Kawashima et al., 2016) is sufficient to capture the transformation from stimulus to activity (Figure 5A_1_-A_2_).

Our previous analysis indicated that the most important transformation occurs in the Rh 5/6 region of the hindbrain. To explain the observed response types in this region in terms of trigeminal neuron activity we employed the same approach, a linear combination of trigeminal activity followed by convolution with a filter and an output nonlinearity (Figure 5B; Figure S5 C-G). Two cell types in this hindbrain region, Slow-ON and Slow-OFF, were similar to the trigeminal cell-types. The Slow-ON type has slightly faster dynamics than the trigeminal ON cells, as evidenced by the adaptive component in its filter, and it is almost exclusively driven by excitatory inputs from the trigeminal ON cells (Figure 5B_1_). We note that since adaptive filters compute a derivative of their input they are prone to increase noise present in their input. The long timescales observed in the filters could therefore reflect a mixture of suppressing noise in the best fit (see Figure S5H-I for a simulation) as well as true neuronal processes underlying adaptation and bursting (Bean, 2007; Vilin and Ruben, 2001). The Slow-OFF type is an almost direct copy of trigeminal activity, evidenced by a filter that is essentially a delta function (Figure 5B_2_). The three other cell types in Rh5/6 critically relied on inhibitory inputs (Figure 5B_3_-B_5_). While trigeminal neurons express a variety of neurotransmitters, trigeminal fibers are largely glutamatergic (Lazarov, 2002). Since rhombomeres 5 and 6 on the other hand contain both inhibitory and excitatory neurons (Kinkhabwala et al., 2011) we used the Slow-ON and Slow-OFF types in the model to provide the required inhibition instead of relying on the trigeminal inputs directly. Both the newly arising Fast-ON and Fast-OFF types required inhibition by Slow-OFF cells, however to differing degrees. Furthermore, as suggested from their activity profiles both their filters signal strong adaptation (Figure 5B_3_ and 5B_4_). The last activity type, termed “Delayed-OFF” as it had fast kinetics but started to respond after the Fast-OFF cells, relied on inhibition by the Slow-ON type. The linear filter of this type potentially hints at a mixture of integration and differentiation, which would also be suggested by the activity profile itself (Figure 5B_5_). To test the importance of temporal filtering and hence the dynamic structure of our model, we fit an alternative model in which no linear filters were applied for cell types in rhombomeres 5/6. As expected such a model still explains the activity of Slow-ON and ‐OFF types well whereas the activity of the Fast-ON and Fast-OFF types could not be recreated from the trigeminal inputs without filtering (Figure S5J). This indicates that temporal filtering of activity is a critical step in creating important response types observed in Rh 5/6.

Since the identified motor cells in the brain did not track the stimulus itself but much like the behavior had a probabilistic chance of firing depending on stimulus strength, we used a simple linear rate-coding model for the transformation from heat-modulated hindbrain activity to activity rates in the motor cells (Figure 5C). Namely, linear combinations of activity in Rh 5/6 cell types explained the activity of each identified motor cell type (Figure 3). This revealed varying combinations of activating and inhibiting influences consistent with the mix of excitatory and inhibitory neurons in this region (Kinkhabwala et al., 2011; Figure 5C_1_-5C_5_). Most motor cell types are activated by the Slow-ON type with the expected exception of Spontaneous-motor cells which receive strong excitation from Slow-OFF cells almost exclusively (Figure 5C_5_). This paradoxical control of spontaneous behavior by a stimulus driven cell type is necessary to explain their lack of activity during stimulation. Interestingly, “Swim” cells receive their strongest excitation from the Fast-OFF cell type (Figure 5C_3_), consistent with the requirement of this cell type in explaining swimming behavior via a simple linear model (Figure 4B”).

The last stage of the model linearly links motor cell output to the observed swim and flick behavior (Figure 5D). Both of these behaviors are strongly activated by the All-motor and Evoked-motor cells. Flick cells overall have a weak contribution to both behaviors but as expected while they inhibit swims they activate flicks. Swim cells on the other hand have a strong contribution to swims but do not influence flicks (Figure 5D).

In summary, we could derive a realistic dynamic circuit model that can explain the observed transformations in activity from sensation to behavioral output and which makes clear and testable predictions about the underlying circuit.

### The circuit model can predict behavior and neuronal responses to novel stimuli

The dynamic circuit model of the sensori-motor transformations (Figure 6A) was created such that each stage of the model was fit independently. This means that errors could accumulate across the model preventing prediction of behavioral output in response to sensory input. Therefore, as a first test of the model, we tried to predict the behavioral output of the experiments that were used to fit the model given the sensory input. Both the rates of swims and flicks were very well predicted by the circuit model (Figure 6B), indicating that errors do not accumulate across the different stages.

**Figure 6.**
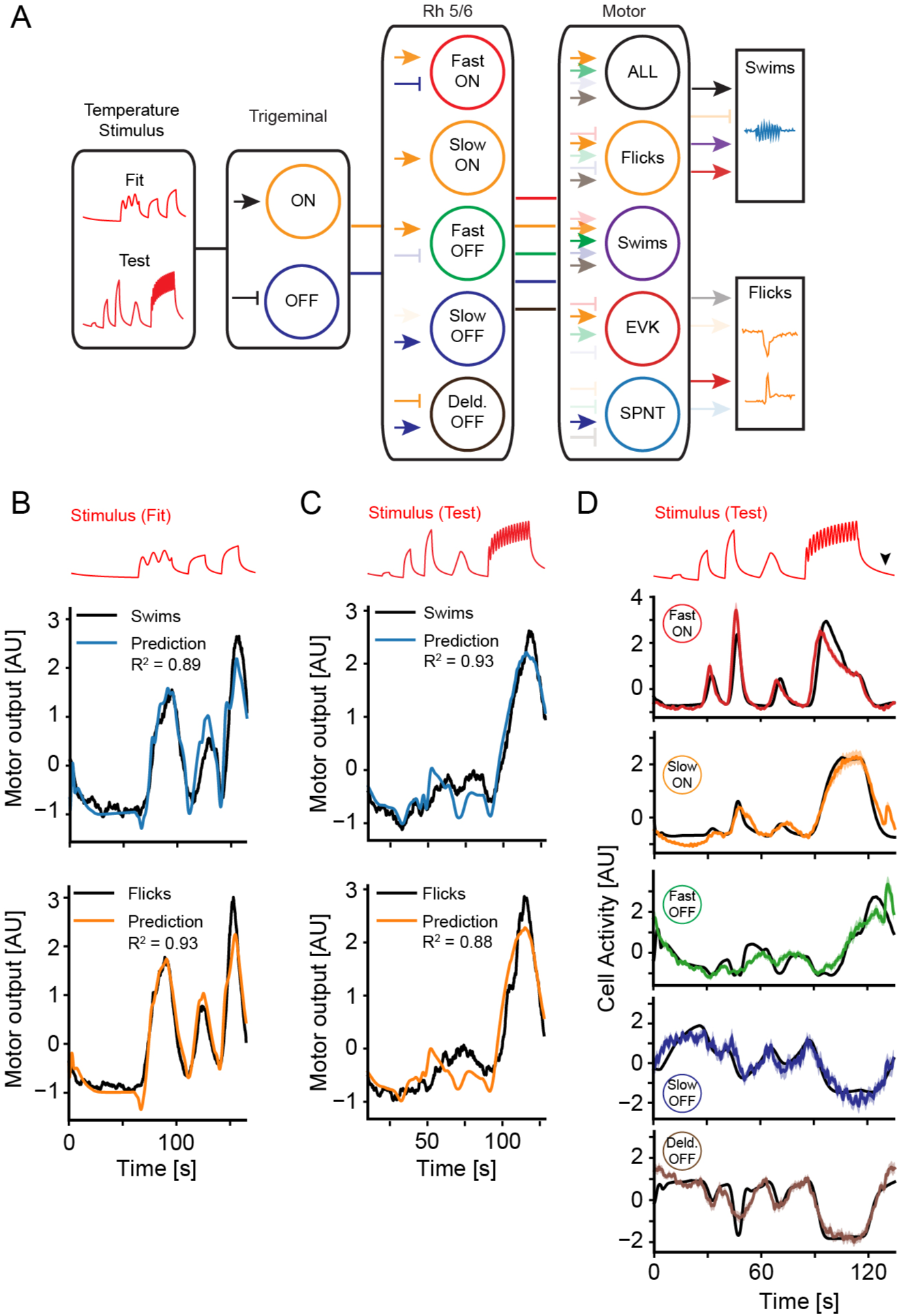
The model predicts behavioral and neural activity in response to novel stimuli. A) Schematic of the full feed-forward model. Colored arrows depict the mixing of sensory input or activity in a previous stage with arrowheads indicating positive (activating) and bars indicating negative (inhibiting) effects. Opacity of arrows indicates the weight of a given component in the fit. B) Prediction of swims (top) and flicks (bottom) based on the model for the same experiments that were used to fit the model. Colored lines represent prediction, black line is observed behavior convolved with the calcium kernel. Stimulus depicted on top for reference. C) Prediction of swims (top) and flicks (bottom) based on the sensory input delivered and motor output observed during the heat and tap experiments. Note that periods in which the behavior is affected by the tap itself have been excluded from the plot. Colored lines represent prediction, black line is observed behavior convolved with the calcium kernel. Stimulus depicted on top for reference. D) Cluster average activity (colored lines) of the indicated types versus each model predicted regressor (black line). Stimulus is depicted on top for reference. The black arrowhead indicates timing of the tap which was present in this test stimulus but which was not included in our modeling. Shading indicates bootstrap standard error. See also Figure S6.

To test the generality of the model, we wanted to test its prediction in response to a sensory stimulus not used for fitting. To this end we used a heat stimulus with distinctly different temporal dynamics consisting of three temperature steps and a ramp followed by a faster oscillating sine wave (Figure 6A and Figure S6A). This stimulus served as a test input to our circuit model to compare the prediction of flick and swim behavior to the actual behavioral rates produced by fish during those experiments. The model did very well in predicting behavior to this novel stimulus, explaining close to 90% of the variance in both flick and swim behaviors (Figure 6C). This indicates that the model does generalize across stimuli of different dynamics.

Encouraged by these results we were wondering whether it might be possible to identify the different heat response types present in Rh 5/6 in this experimental set using model predictions as regressors. Probing activity using the model predictions indeed recovered cells with correlated activity for each predictor (Figure S6B). Clustering cells by correlation into types, recovered type average activity that matched the individual model predictions to a large extent as evidenced by the close juxtaposition of the predicted activity and cluster average activity (Figure 6D). Importantly the average activity profiles of the types matched expectations. This can be seen by comparing average activity of Fast-ON with Slow-ON types, where, as in the other experiment, Fast-ON cells showed quicker onset responses followed by adaptation compared to a more sustained profile in the slow type (Figure 6D). The test stimulus set also uncovered that Fast-OFF cells do not only increase their activity on temperature decline but seem to be especially inhibited during temperature rises. This is revealed by the fact that activity in this cell type started to increase before offset of the sine stimulus but concomitant with a decrease in Fast-ON activity (around 100 s into to the trial instead of 115 s, Figure 6D).

Interestingly, the prediction of swims and flicks depends to a different extent on accurate representation of Rh 5/6 activity. Using the model without temporal filtering (Figure S5J), prediction of flicks is nearly as good as for the full model, while the prediction of swims is considerably worse for both stimuli, as evidenced by a clear drop in explained variance (Figure S6C-D). This is in line with the ability to predict flicks purely based on trigeminal activity (Figure 4A”). As expected, this comparison model performs much worse in identifying Fast-ON and Fast-OFF cells in Rh 5/6 in response to the test stimulus compared with the full model (Figure S6E).

In summary, we could demonstrate that our activity constrained circuit model generalizes to novel stimuli and is able to predict both behavioral output and intermediate neuronal activity in this context. This argues that the model accurately represents computations during sensori-motor transformations in heat perception.

## Discussion

The sense of temperature is conserved from mammals (Murakami and Kinoshita, 1977) to bacteria (Maeda et al., 1976) but how information about environmental temperature is represented across the brain, especially in vertebrates, is largely unknown. In this study, we combined a novel experimental paradigm with theoretical modeling approaches to delineate the circuits and computations that underlie the transformation from temperature sensation to behavior. Performing 2-photon functional calcium imaging with subsequent volumetric image registration allowed constructing a comprehensive atlas of heat processing centers in the larval zebrafish brain. Furthermore, the combination of functional recordings during heat stimulation and behavioral readout revealed critical response transformations in the hindbrain that are necessary to explain the observed behavior. These transformations change a sensory representation that is largely confined to reporting heat levels to a rich representation that extracts features such as direction of temperature change. To formalize these transformations, we built a dynamic circuit model constrained by anatomy and the observed activity. By concurrently fitting connectivity strength and “filter kernels” we created a framework that allows predicting behavior and neuronal activity in response to diverse heat stimuli.

### A brain wide atlas of temperature modulated activity

Previous work across animal species has provided important insight into the cellular and molecular mechanisms and circuit logic of temperature detection, however brain wide analysis of temperature modulated activity is lacking. Here, we mapped neurons with heat modulated activity across a whole vertebrate brain and found that the representation of temperature is widespread and especially prominent in the fore‐ and hindbrain. As expected the trigeminal ganglion, a somatosensory area, contains heat sensitive neurons. However, only a small fraction of cells in the trigeminal ganglion responded to our heat stimuli (Figure 2E), which were well outside the noxious range (Figure 1B). This is in line with previous reports in mice where only few trigeminal neurons detect innocuous warmth while many respond to noxious heat (Yarmolinsky et al., 2016). The rhombomere 5/6 region, which is a target region of the trigeminal, contained prominent clusters of heat modulated cells. Furthermore, responsive cells were aggregated in the dorsal cerebellum, in both habenulae as well as in pallium and sub-pallium. This likely indicates that temperature not only influences behavior directly but also provides meaningful information for higher order processes controlled by the brain. Heat modulated activity in the pre-optic area hints at a potential functional conservation of this structure which is involved in the regulation of body temperature in mammals (Boulant, 2000) and involved in behavioral fever in toads (Bicego and Branco, 2002).

### Functional diversity of temperature encoding

Across the brain we find that temperature is encoded by both ON and OFF type cells. A separation of temperature coding into ON and OFF cells also emerged as a common principle from previous studies in flies (Frank et al., 2015; Liu et al., 2015) and mice (Ran et al., 2016). Such a separation of sensory coding into ON and OFF channels has been thought to aide in coding efficiency (Gjorgjieva et al., 2014) and may serve to mitigate effects of correlated noise. Comparing heat responsive cell types across regions revealed clear differences in response dynamics. Sensory neurons in the trigeminal fall into two strongly anticorrelated ON and OFF types with sustained responses. In the hindbrain on the other hand more divergent response types could be identified including cells that showed strong adaptation and that are therefore most sensitive to changes in temperature. While we observe a combination of sustained and transient ON and OFF cells in Rh 5/6 (Figure 4B), second order neurons in the mouse spinal cord represent warming exclusively with sustained responses and cooling in a transient manner (Ran et al., 2016). The forebrain, especially the pallium and pre-optic area, harbored cell types that seem to represent temperature on slower timescales (Figure S4 B-E). This could indicate that these cells set long-term states rather than control behavior itself.

### Stimulus and behavior separate motor activity across the brain

Using correlational analysis, we mapped motor related activity across the brain. Compared with the stimulus representation, motor related cells are considerably more localized. A large fraction of motor cells is localized in the anterior hindbrain as well as in the cerebellum, slightly ventral of the stimulus related cell clusters. On the other hand, only a small fraction of cells in the forebrain displayed motor-correlated activity. A simple criterion based on tail dynamics allowed subdividing the head-embedded behavior into two broad classes, “swims” and “flicks”. Swims represent bouts which likely correspond to straight swims and routine turns in freely swimming behavior, while flicks are unilateral tail deflections that likely correspond to strong, in-place turns. Mirroring the difference in behavioral output, we find separate pools of cells that are correlated with swims and flicks indicating that at the level of the hindbrain separate pools of cells are used to initiate these different behaviors. This is similar to the subdivisions of reticulo-spinal neurons for straight swim or turn initiation (Huang et al., 2013; Orger et al., 2008). In line with ipsilateral cells controlling directional turns (Orger et al., 2008), cells encoding left flicks are more prominent in the left hemisphere of the hindbrain and vice versa (70% of cells are ipsilateral, Figure 3F).

We also identified stimulus specific motor cells. “Evoked-motor” cells are activated almost exclusively during bouts while the stimulus laser is on, while “Spontaneous-motor” cells are mostly active during spontaneous bouts in the absence of our heat stimulus (Figure S3B). This separation might underlie temperature induced changes in bout structure such as observed increases in average bout speed caused by temperature increases in freely swimming larval zebrafish (Haesemeyer et al., 2015). That spontaneous and evoked behaviors are controlled by separate pools of cells is in contrast to Aplysia, where changes in distributed activity in the same pool of cells accounts for differences in spontaneous and evoked behaviors (Wu et al., 1994). Previous studies in zebrafish however support a coding strategy whereby different behavioral modules are controlled by separate pools of cells (Orger et al., 2008; Thiele et al., 2014).

### A dynamic circuit model of sensori-motor transformations during heat perception

Models constrained by behavioral and physiological data provide important insight for understanding the logic of sensori-motor transformations (Clark et al., 2013). Recently, modeling approaches have been used in the larval zebrafish to understand processes underlying prey-selection (Bianco and Engert, 2015) as well as the generation of the optomotor response (Naumann et al., 2016). This approach resulted in a realistic multi-scale circuit model of optomotor induced turning revealing the circuit logic of binocular stimulus integration (Naumann et al., 2016). We similarly used known anatomy and observed neural activity to derive a realistic circuit model of temperature perception. Correlational analysis strongly suggested that a critical step in the sensorimotor transformations is the observed change of temperature representation between the trigeminal ganglia neurons and their rhombomere 5/6 target region in the hindbrain. This transformation especially improves the prediction of undulating swim behavior while flicks are already fully predicted at the level of sensory activity in the trigeminal ganglia. We therefore reasoned that the simplest plausible circuit would consist of the trigeminal neurons, the cells in their Rh 5/6 target area as well as the identified motor cells. Since the stimulus space is characterized by the dynamics of temperature change, we had to extend the previous modeling approaches and devise a circuit model that takes these dynamics into account. In fact, a comparison model using static rate coding alone fails to replicate activity and behavior (Figure S5J and S6C-E). For stimulus encoding and the transformation of temperature representation in the brain, the model therefore consists not only of linear coefficients but also filter kernels that are fit for each cell type separately. These filter kernels allow quantifying the dynamical changes in heat representation (Figure 5 A-B). Using this approach, we find that while Slow-OFF cells in Rh5/6 largely copy their trigeminal input, other cell types such as the Fast-ON and ‐OFF cells rely on adaptation to transform their inputs.

In the model, the filter kernels are a property of a given cell type, effectively acting as input filters. This is a plausible explanation as different cell intrinsic processes can lead to the observed spiking adaptation in the Fast-ON or ‐OFF cell type or more complex interactions between adaptation and bursting behavior as suggested in the filter of the Delayed-OFF type (Blair and Bean, 2003; Friedman et al., 1992; Kernell and Monster, 1982; Pedarzani and Storm, 1993). Further experiments using patch clamp recording in identified cells are needed to decide whether these are indeed properties of the cells or rather emerging features of local circuits. Furthermore, the model suggests long timescales on the filter properties. These timescales are plausible for processes such as late adaptation observed in motoneurons or hippocampal neurons (Kernell and Monster, 1982; Pedarzani and Storm, 1993) or slow afterpotentials involved in cortical bursting (Bean, 2007; Friedman et al., 1992). On the other hand, the nature of calcium imaging together with the inherent noise precludes a full quantitative interpretation of the filter kernel timescales (FigureS5H-I). They do however, make clear qualitative statements about the expected cellular properties which can be confirmed with future experiments using electrophysiological recordings or voltage imaging.

### The circuit model as a framework for hypothesis testing

Our circuit model makes clear and testable predictions about the computations and architecture underlying the sensori-motor transformations during heat perception. While trigeminal fibers are largely glutamatergic (Lazarov, 2002), three of the five cell types identified in rhombomeres 5 and 6 in the hindbrain rely on inhibitory inputs. The model therefore posits that part of the Slow-ON and Slow-OFF cell-types should be inhibitory interneurons. At the same time, excitatory projections from these neurons onto Motor cells are required to explain the activity rates of some Motor types. This possibility is well supported by previous anatomical studies that identified both glutamatergic and glycinergic neurons in this region (Kinkhabwala et al., 2011).

Our previous behavioral study predicted that straight swims should be activated by a strong OFF signal which is less influential for turning (Haesemeyer et al., 2015). This study supports this conclusion: the circuit model predicts that swim motor cells are most strongly driven by the Fast-OFF cell type (Figure 5C_3_).

The core transformation from the trigeminal to the hindbrain to the motor output is well captured by our dynamic circuit model which is constrained by a variety of activity measurements and which we validated by testing its predictive power against novel sensory stimuli. Such a brain wide realistic model that captures the dynamic aspects of sensorimotor transformations is novel in the context of temperature processing and provides a computational and experimental framework for generating testable circuit models of temporal coding. Furthermore, the general model architecture presented here will allow to include modules responsible for higher order processing, such as observed activity in the cerebellum and forebrain areas, in the future and can be easily applied to other stimuli and organisms to capture similar transformations in representation.

## Materials & Methods

All experiments were conducted on 6-7 days post fertilization zebrafish of the strains indicated below. Fish were fed paramecia from day 5 onwards. All experiments followed the guidelines of the National Institutes of Health and were approved by the Standing Committee on the Use of Animals in Research of Harvard University. All analysis was performed using software custom written in Python 3.5.

### Imaging and behavior

All experiments used for mapping heat responsive neurons and for model derivation used nuclear expressing Huc-H2B-GCaMP6s fish (Freeman et al., 2014). Experiments combining heat and taps were performed in cytoplasmic Huc-Gcamp6s fish (Wee et al., in preparation).

Larval zebrafish were embedded in 2.5 % medium melt agarose (Fisher scientific, USA) and their tails were freed the night before the experiment. Experiments were conducted in a custom built 2-photon microscope and run using custom written software in C# (Microsoft, USA). Heat stimuli were delivered using a 1 W 980 nm fiber-coupled diode laser (Roithner, Austria) coupled into a collimator (Aistana Inc., USA) placed under the microscope objective 4 mm in front of and 1.2 mm above the head of the zebrafish larva pointing downwards at an angle of 16.5 degrees. The laser power was controlled by the computer via a laser diode driver (Thorlabs, USA). We note that the 980 nm laser itself did not excite GCaMP fluorescence due to the low photon density. The main mapping experiments consisted of the imaging of 30 individual planes, spaced 2.5 *μ*m apart. In each plane 3 trials of the stimulus depicted in Figure 1B were presented.

The heat and tap experiments consisted of imaging 4 individual planes, spaced 5 *μ*m apart. In each plane 25 trials of the stimulus depicted in Figure S1G were presented.

To avoid excessive heating of the preparation by scanning over the eyes, custom exclusion masks were created for each experiment in which the eyes were in the field of view restricting the scan-lines such that the eyes were excluded from the field of view.

### Image stabilization

To counter heat-induced deformations of the preparation, before each plane was scanned a +/- 5 *μ*m sized pre-stack consisting of 21 slices spaced 0.4 *μ*m apart was acquired. During scanning, each acquired plane was crosscorrelated with each plane in the pre-stack and using a low-pass filter the position of the objective was adjusted online so as to minimize z-drift. Since the heat induced drift observed in RFP stacks followed very reproducible kinetics these were used to predict the movement induced by heating during our experiments to induce very slight movements in the predicted direction. This served to overcome the delay induced by our low-pass filter.

### Segmentation

Activity traces were obtained from all experiments using anatomical segmentation. Nuclear GCaMP stacks were segmented anatomically using Cell Profiler (Carpenter et al., 2006). To segment the cytoplasmic GCaMP stacks we used the nuclear exclusion of the indicator to define cell-centroids based on a minimum filter. This was followed by growing a mask using temporal correlation of individual pixel timeseries up to an anatomically defined size.

### Registration and annotation

3D image registration based on CMTK (Rohlfing and Maurer, 2003) was used to create a nuclear GCaMP-6s reference stack to which all experimental stacks were registered as described elsewhere (Portugues et al., 2014; Randlett et al., 2015). The reference brain was used to annotate anatomical regions of interest based on Z-Brain annotations (Randlett et al., 2015).

### Clustering of heat and motor related activity

To identify motor related activity, motor regressors were created by convolving a bout start trace with an exponential calcium kernel with a decay half time of 3 s. This decay time was derived from motor triggered averages across hindbrain neurons. Behavioral subtype regressors were created by only considering bouts of a given type. The behavioral regressors that differentiate stimulus and rest periods were created by only considering motor events during those respective phases. Every cell with a correlation of at least 0.6 to at least one motor regressor was considered a “motor cell”. Since not all behaviors were observed in all imaging planes, cells were only assigned to a more specific category (such as swims versus all bouts) if the correlation to the more specific regressor was significantly higher (*p* < 0.01, bootstrap hypothesis test) than to any more general regressor.

To identify sensory response types, all motor correlated cells were first removed from the data using a lower correlation cutoff of *r* > 0.4. Subsequently, to identify ON and OFF cells across the whole brain (Figures 1 and 2) filtering criteria based on stimulus modulation and a requirement of correlated partners across cells were used to remove around 90 % of cells from further consideration for computational purposes (see Supplemental Materials and Methods for details). For region-specific clustering of activity (Figure 4) each nuclear centroid was assigned to an annotated region. For this data the filtering step was omitted as the datasets were of manageable size already. For each whole-brain or region-specific set cell-to-cell correlations were used to build a similarity graph. Subsequently spectral embedding followed by K-means clustering was used to calculate minimum cuts in this graph thereby naturally separating the data into clusters of highly correlated activity. To further prune the clusters, the cluster average activity was used as sensory regressors and only those cells were kept as cluster members that showed a correlation of at least 0.6 to the cluster average. We defined the number of clusters such that at least one empty cluster was obtained (i.e. a cluster with average activity to which no other cell was highly correlated).

To calculate ΔF/F0 values for reporting cell fluorescence we used the average across the first baseline period as the resting fluorescence F0.

### Modeling

The general structure of our feed-forward model relied on two “dynamic” stages, the encoding of the sensory stimulus in the trigeminal ganglion as well as the transformation of trigeminal activity in Rh5/6. These were followed by two rate-coding steps, the transformation of firing rates in Rh5/6 into firing rates of the motor cells as well as the transformation of motor cell rates into behavioral rates (Figure 5). In each stage, linear coefficients were fit that represent the activations of cells (or behavior) by the different cells (or the stimulus) in the previous stage. To allow for changes in the dynamical representation, in addition convolutional filter kernels were fit in the first two stages of our model. Since we expected stimulus encoding to be mostly governed by the nuclear GCaMP6s calcium kernel the filter of the trigeminal ganglion was parametrized with an on (*τ*_*ON*_) and an off-rate (*τ*_*OFF*_) according to:

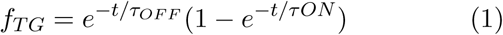

Since the differences in activity profiles between trigeminal neurons and neurons in Rh 5/6 suggested that some cells performed a differentiation of their input a filter parametrization that allows for both positive and negative (adapting) components was used. The filter was parametrized by a scaling term s and two rates *τ*_1_ and *τ*_2_ according to:

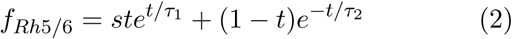

The filter parameters were derived together with the linear coefficients using Markov-chain Monte Carlo (MCMC) sampling via PyMC3 (Salvatier et al., 2016).

Since the relationship of predicted to actual values after fitting suggested a nonlinear transformation, for the two first stages of the model a cubic nonlinearity of the form

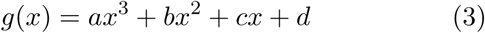

was fit as well using least squares optimization.

For the last two rate-coding steps the linear coefficients were fit using MCMC as well and no nonlinearities were necessary. Reported confidence intervals in all cases are 99 % confidence intervals based on draws from the approximated posterior distribution. See supplemental materials and methods for details of model definitions and prior parameter distributions.

## Author Contributions

MH conceived the project in discussion with FE and AFS, and carried out all experiments. DNR and JML built the 2-photon microscope and wrote the imaging software. MH wrote the online stabilization pipeline and all analysis software, analyzed the data and built the model. MH, AFS and FE interpreted the data and wrote the manuscript.

## Acknowledgements

MH was supported in part of this project by an EMBO Long Term Postdoctoral fellowship (ALTF 1056-10) and a postdoctoral fellowship by the Jane Coffin Childs Fund for Biomedical Research. Research was funded by NIH grants DP1-NS082121, U01NS090449 and 5R24NS086601 to FE. We thank Ruben Portugues, Andrew D. Bolton, James Fitzgerald and Aravi Samuel for critical discussion and helpful comments on the manuscript.

